# Quantifying tumor specificity using Bayesian probabilistic modeling for drug target discovery and prioritization

**DOI:** 10.1101/2023.03.03.530994

**Authors:** Guangyuan Li, Anukana Bhattacharjee, Nathan Salomonis

## Abstract

In diseases such as cancer, the design of new therapeutic strategies requires extensive, costly, and unfortunately sometimes deadly testing to reveal life threatening “off target” effects. A crucial first step in predicting toxicity are analyses of normal RNA and protein tissue expression, which are now possible using comprehensive molecular tissue atlases. However, no standardized approaches exist for target prioritization, which instead rely on ad-hoc thresholds and manual inspection. Such issues are compounded, given that genomic and proteomic data detection sensitivity and accuracy are often problematic. Thus, quantifiable probabilistic scores for tumor specificity that address these challenges could enable the creation of new predictive models for combinatorial drug design and correlative analyses. Here, we propose a Bayesian Tumor Specificity (BayesTS) score that can naturally account for multiple independent forms of molecular evidence derving from both RNA-Seq and protein expression while preserving the uncertainty of the inference. We applied BayesTS to 24,905 human protein-coding genes across 3,644 normal samples (GTEx and TCGA) spanning 63 tissues. These analyses demonstrate the ability of BayesTS to accurately incorporate protein, RNA and tissue distribution evidence, while effectively capturing the uncertainty of these inferences. This approach prioritized well-established drug targets, while deemphasizing those which were later found to induce toxicity. BayesTS allows for the adjustment of tissue importance weights for tissues of interest, such as reproductive and physiologically dispensable tissues (e.g., tonsil, appendix), enabling clinically translatable prioritizations. Our results show that BayesTS can facilitate novel drug target discovery and can be easily generalized to unconventional molecular targets, such as splicing neoantigens. We provide the code and inferred tumor specificity predictions as a database available online (https://github.com/frankligy/BayesTS). We envision that the widespread adoption of BayesTS will facilitate improved target prioritization for oncology drug development, ultimately leading to the discovery of more effective and safer drugs.

**Code and Data Availability:** The BayesTS code and inferred tumor specificities are available at (https://github.com/frankligy/BayesTS), along with the scripts for reproducing the results in the study and instructions on retraining the model. The training data is available at Synapse (https://www.synapse.org/#!Synapse:syn51170082/files/).

## INTRODUCTION

Tumor cells undergo extensive genetic and genomic alterations that substantially alter their cell growth and metabolism ^1,2^. While advantageous for tumor survival, these alterations also introduce potential molecular vulnerabilities that can be exploited for the design of new targeted therapies. For example, Imatinib, a Tyrosine Kinase Inhibitor, is an effective targeted cancer therapy that acts by selectively binding with the BCR-ABL1 fusion protein in Chronic Myelogenous Leukemia (CML) patients, substantially improving clinical outcome ^3,4,5^. New and existing cancer immunotherapies, such as checkpoint inhibitors and adoptive cell transfer have significantly increased the repertoire of targets for cancer therapy by leveraging a patient’s own immune system as “living drugs” ^6,7^.

While promising clinical results have been reported in multiple malignancies ^8–11^, the efficacy of both targeted and immunotherapy approaches depends on the improved selection of molecular targets, ranging from cancer surface proteins, HLA presented neoantigens and novel tumor-specific epitopes ^12^. Target specificity is frequently best assessed on the basis of target gene or protein expression in diseased versus non-diseased tissues. Indeed, selective evaluation of non-diseased tissues alone can inform tumor specificity prediction. Importantly, “on-target off-tumor” toxicity can result in severe side effects which can result in significant morbidity and mortality when a therapy is evaluated in clinical trials ^13,14^. For example, neurotoxicity is a known side-effect of Chimeric antigen Receptor T cell (CAR-T) therapy, with a potential mechanism identified through single-cell genomics of brain mural cells, which express the CAR-T target, CD19^15^.

Quantifying the tumor specificity of drug targets is a non-trivial task, as existing molecular measurement data is heterogeneous, with the need for multimodal (e.g., RNA, protein) measurements. Gene expression can be quantified using multiple counts-based approaches, with variable numbers of samples per tissue assayed for and variable molecular distributions. Further, it is well documented that mRNA expression levels do not often correlate with their protein abundance, which is subject to extensive post-transcriptional and post-translational regulation ^16^. Hence, relying solely on one single molecular modality and quantification approach is problematic when gauging target specificity. Current best practices for identifying tumor-specific targets rely on the application of an array of fixed or tunable thresholds ^13,17^, which can be useful but also introduce subjectivity and substantial variance across studies and sample cohorts. Such ad-hoc methods further hinder the analysis and exploration of target specificity. For example, if a transcript is detected at low levels in some samples due to shallow sequencing, but is detectable through alternative methods, a fixed threshold may incorrectly classify the gene as non-specific. Similarly, if the distribution of gene expression is highly skewed, a fixed threshold may not accurately capture the underlying biology. While recent neural network based methods could be applied to learn such complex nonlinear relationships, such models are not sufficiently interpretable and do not account for uncertainty to be explicitly quantified. In contrast, new Bayesian probabilistic approaches present an intriguing potential solution to this challenge, as they are able to incorporate prior knowledge and uncertainty in a principled manner, leading to accurate and interpretable predictions. Such modeling in single-cell genomics ^18,19^, compositional analysis ^20^ and spatial deconvolution ^21^, has proven valuable, where missing and heterogenous molecular measurements are inherent.

Here we describe the first tunable and quantitative tumor specificity score termed BayesTS, using probabilistic modeling. Bayesian probabilistic modeling is well-suited to quantify target specificity given prior knowledge of the distribution of multimodal measurement data which can be incorporated into a single coherent model. This approach was designed to handle missing data and to explicitly quantify uncertainty in any supplied molecular measurement data. We demonstrate that BayesTS can successfully learn from multiple molecular modalities and can differentiate between proven safe versus high-risk CAR-T therapy targets. We show that BayesTS enables accurate dynamic tuning of tissue importance, and can identify novel tumor specific targets. This approach can further be generalized to new modalities aside from gene products, such as splicing neoantigens, which represent new promising targets for therapy. Our proposed Bayesian Tumor Specificity (BayesTS) score is efficient and scalable, using variational inference and leveraging the power of the PyTorch framework to run on GPUs ^22,23^ with effective subsampling. These features enable BayesTS to compute tumor specificity scores for large datasets of 100,000 targets in less than 1 minute.

## METHODS

### BayesTS model

BayesTS was developed as a hierarchical Bayesian model that jointly considers multiple types of molecular measurements that independently quantifies tumor specificity for a target. In our model, we consider three separate observations, namely, tissue distribution of the bulk RNA-Seq counts (**X**), normalized bulk RNA-Seq counts (**Y**) and protein-level expression derived from immunohistochemistry slides (**Z**). The provided base model can potentially be extended to incorporate other modalities (e.g. single-cell cell-type distributions). We define the tumor specificity as the expression of a target in normal tissues bounded by 0 and 1, where a smaller value should represent a more confident potential target (e.g., tumor specific) where a higher value represents greater risk of being non-specific (**Figure 1A**). Non-diseased tissue bulk RNA-Seq data was obtained from GTEx and TCGA matched controls (see Bulk RNA-Seq preprocessing), and the normal tissue protein immunohistochemistry annotations were downloaded from Human Protein Atlas (HPA) ^24^ (see Immunohistochemistry preprocessing).

**Figure 1.**
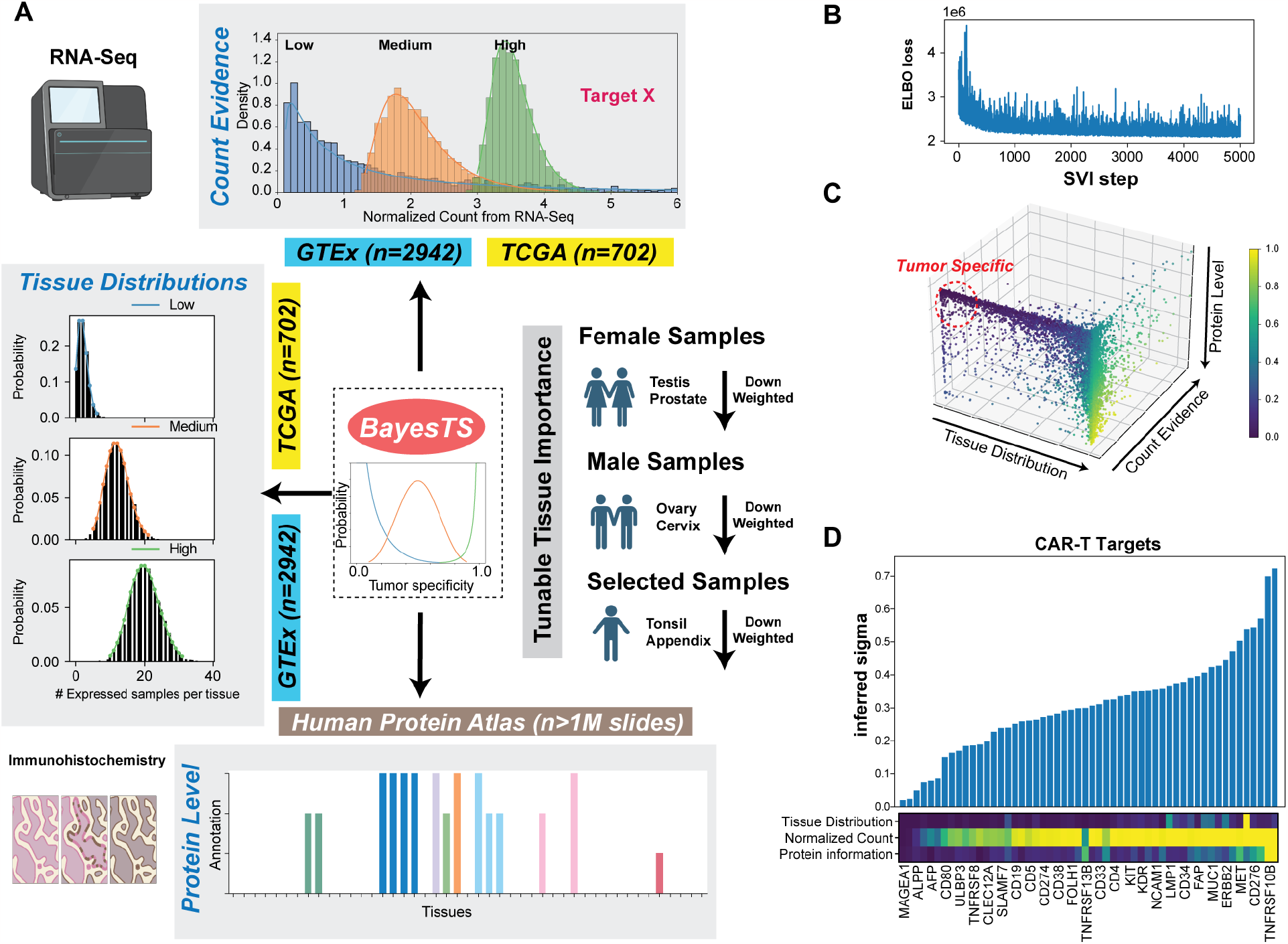
BayesTS automates the identification of tumor specific proteins from multiple data dimensions. A) Schema of the BayesTS model. A single continuous value (BayesTS) bounded by 0 and 1 gives rise to three domains of evidence including the normalized RNA count values across multiple samples (top), the number of samples expressing a specific target (left), and immunohistochemistry evidence indicating whether a protein is confidently detected (bottom). Tunable user-supplied tissue importance options enable flexible modeling of tumor specificity (right). B) The ELBO loss of the BayesTS model with training step shown in x-axis. C) Inferred tumor specificity of 13,306 protein coding genes (with available protein information) across the three evidence domains (axes). D) Inferred tumor specificity of 55 CAR-T targets in at least one current clinical trial (subset of genes shown). (Bottom) Heatmap of the ground-state evidence for each.

Here, we denote the expression level in normal tissues (tumor specificity) as σ and assume all observations are generated by this underlying latent variable through a hierarchical sampling process. Specifically, the shape and values of the three observations are explained below. A tissue distribution observation **X**_**it**_, where **i** denotes a given target and **t** denotes a tissue type, **i** can be any druggable target (e.g., gene, protein, splice junction, MHC-presented neoantigen, mutation, etc). In our default model, the total number of tissues considered is **T**, where each value in the matrix **X** represents the percentage of samples in each tissue type in which this target can be detected. A Normalized bulk RNA-Seq observation **Y**_**is**_, where **s** denotes a normal sample in our model and the total number of samples considered is **S** and each value in matrix Y represents the normalized count value (Count Per Million (CPM)) for each target in the sample. Last but not least, the protein annotation observation is defined as **Z**_**ip**_, where **p** denotes the number of immunohistochemistry slides for a target. In our model, the total number of slides considered is **Z**. Each value in matrix **Z** represents a categorical value {0, 1, 2, 3} which corresponds to the HPA consortium annotations {high, medium, low, not detected}. The plate notation for this hierarchical model is shown in **Supplemental Figure 1**.

**Supplementary Figure 1.**
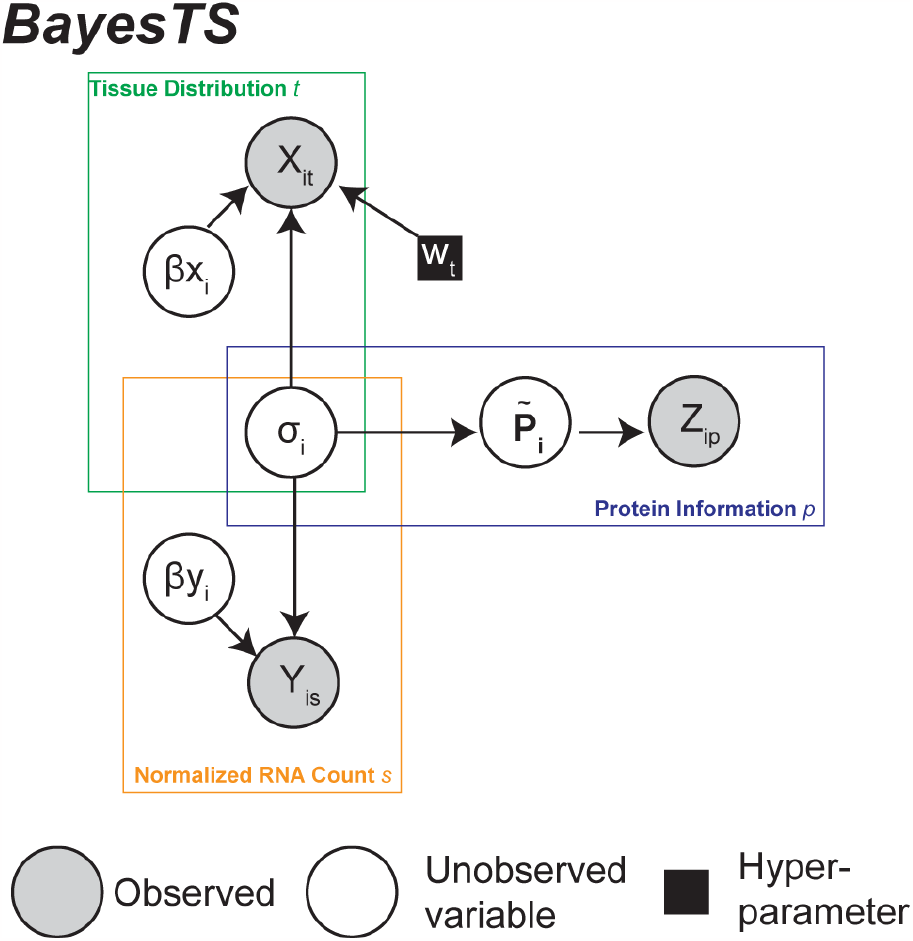
The plate notation of the BayesTS model. Each circle represents a random variable in the model, shade denotes the observed values and hollow denotes the unobserved values. Black squares represent tunable hyper-parameters.

Mathematically, the model can be described as the following. First, since the inferred tumor specificity (σ) is strictly bounded by 0 and 1, we use a Beta distribution to represent this latent variable with a weakly informative prior Beta(2,2) centered at 0.5.

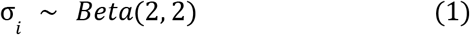

The normalized count value (**Y**) is non-negative and is modeled by a LogNormal distribution with a fixed scale parameter 0.5, whereas the loc parameter is defined by the underlying tumor specificity σ. The intuition is if a target is highly expressed (tumor specificity σ skewed toward 1) then it is highly likely to have a higher mean value for the normalized count value. We use a fixed variance because we do not observe overdispersion and it is also advisable to begin with a low variance model for better training convergence. Another property of LogNormal distribution is the magnitude of values increasing exponentially, which coincide with the normalized RNA count data. We introduce a scale factor β*y*_*i*_ to compensate the different magnitudes between σ and the observed value in **Y**. We use an informative prior Gamma(10,1) to penalize values deviating from its center because value 10 can generate data close to the observed matrix in our simulation experiments.

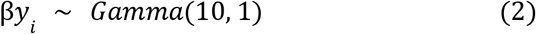

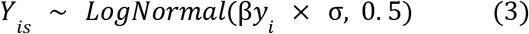

The tissue distribution is to describe RNA expression across different tissues. The input **X** to the model represents the percentage of samples expressing this target with each tissue. Most of the curated tissue types in our reference set have a minimum number of 25 samples (sampled GTEx dataset of matching donors) but increase to hundreds of samples for some tissues (associated with common cancers), we chose 25 as the total count, to transform the ratio data to count data, so that we can model it using Poisson distribution. Similar to the normalized counts value, we use β*x*_*i*_ to account for the difference in magnitude between σ and the observed value in **X**. Here we use 25 as the mean of the scaling factor, because it assumes a target expressed at a medium level should have ∼12.5 samples (half), expressed in each tissue.

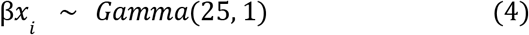

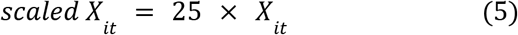

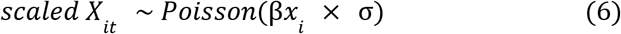

Here we allow the user to tune the tissue importance by introducing a weight vector *w* _*t*_ ∈[0,1], by default, all the values in the tissue weight vector are 0.5, representing equal importance for each tissue. This weight vector is incorporated as the probability parameter in a binomial distribution. We introduce the random variable **Total** that denotes the total number of counts for each tissue:

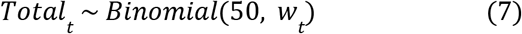

By default, all weights are 0.5, so that the total count for each tissue should be ∼25, which is the same as the vanilla model defined above. However, when users have a strong reason to downweight or upweight certain tissues, *w*_*t*_ becomes important. For example, if we adjust the importance of tonsil to 0.1, then the total count that can be distributed is only around 5 instead of 25 for the tonsil samples. In an extreme case, even if the target is expressed in 90% of tonsil samples, the actual scaled count value the model considers is only 4.5, which won’t affect the model inference as much as it could in default setting.

For the protein data, each entry in the matrix **Z** is assigned a categorical value of 0, 1, 2 and 3 (highly, medium, low and no expression, respectively - see Immunohistochemistry preprocessing). We model the observation **Z** as a categorical distribution where the probability for observing a label as high, medium, low and not detected is derived from the underlying parameter σ. Particularly, when the value of σ is high, the chance of observing the label as high is also increased, and vice versa. Hence, we parameterize the probability vector in the categorical distribution as following:

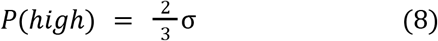

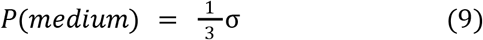

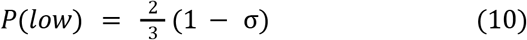

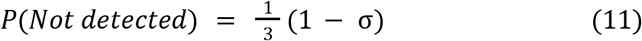

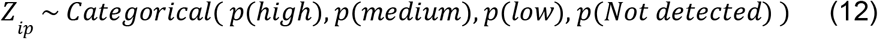

Although the tissue importance is mainly used for tuning the tissue distribution (**Y**), we also populate the weight vector *w*^*t*^ to the protein level to augment the tuning effect (when protein data is available for inference). Here, the weights impact the number of “high” and “not detected” labels in the input **Z** matrix. Specifically, if we denote the original protein expression label distribution of gene i in tissue t as *Z*_*it*_ ∈{0,1,2,3}and the number of high, medium, low and not detected are *n*_0,_*n*_1,_*n*_2,_*n*_3._Given the weight, the updated numbers can be derived as following:

We first derive a scaling factor for each tissuey *sf*_*t*_ based on its ratio against the baseline:

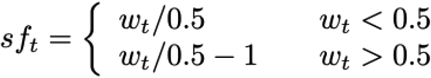

If *w*<0.5:

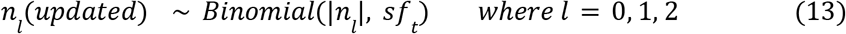

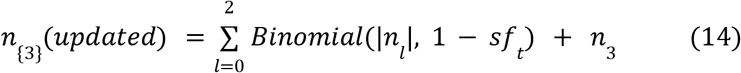

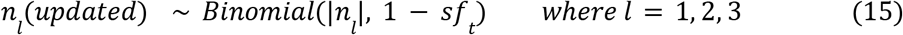

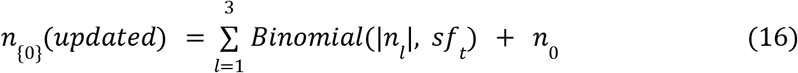

The intuition is, for instance, when the weight of tonsil becomes 0.1, the number of “high”, “medium” and “low” will be reduced by replacing these assignments as “not detected” following in Bernoulli probability equal to the scaling factor 0.2 (0.1/0.5). On the other hand, if the weight is as high as 0.9, then the number of “medium”, “low” and “not detected” will instead be reduced by replacing them as “high” following Bernoulli probability equal to the complement of scaling factor 0.8 (0.9/0.5-1), which arrives at 0.2. When the weight equals the default value of 0.5, no update will be performed.

### Deriving a posterior distribution using variational inference

We derive the posterior distribution for the tumor specificity parameter σ using Variational Inference (VI) as it is faster and more scalable than Monte Carlo Markov Chain (MCMC). Briefly speaking, variational inference aims to construct a simpler distribution (variational distribution)

*q*_ϕ_(σ) that is easy to sample from, to approximate the real posterior distribution *p*_θ_(σ|*x*)after fitting the observed data. It transforms the approximation into an optimization problem as minimizing the differences between two distributions (measured by Kullback-Leibler divergence) is equivalent to maximize the Evidence Lower Bound (ELBO):

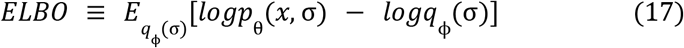

We train the model and maximize the ELBO using Adam optimizer ^25^ with a learning rate set to 0.002 and beta parameters as (0.95, 0.999). Training epoch is set to 5,000 which has been shown to achieve convergence in our experiments (**Figure 1B, Supplementary Figure 2A, 6A**). Since the multiple observations inherently lie on different scales and magnitudes, to avoid the situation where one piece of molecular evidence dominates the training process. We scale the log probability of each distribution based on the empirical evidence by first training each of them. The maximum losses for each single evidence were used to determine the actual scaling factor when combining them (**Supplementary Figure 2A**). We also recommend performing this step when the user extends our model to an internal dataset for retraining, as this will make sure that the derived scaling factor is specific to the datasets being collected.

### Bulk RNA-Seq preprocessing

Raw FASTQ files were downloaded from GTEx following dbGAP authorization using the ANVIL cloud platform. In total 2,942 unique tissue samples corresponding to 52 non-diseased tissues were selected, ranging from 5-360 samples per tissue where 72% samples have around 25 samples. Tissues with less than 10 samples were excluded for tissue distribution analysis due to lack of statistic power. Genome and transcriptome alignment were performed using STAR version 2.4.0h^26^, according to TCGA recommendations, to hg38 and Gencode v36 (https://gdc.cancer.gov/about-data/gdc-data-processing/gdc-reference-files). We strictly follow the guidance of TCGA RNA-Seq manual (https://docs.gdc.cancer.gov/Data/PDF/Data_UG.pdf) as appreciable splice junction differences have been seen in our internal analysis when different STAR versions or genome annotations were used, to jointly analyze GTEx and TCGA normal controls together, in order to minimize possible batch effects. TCGA pre-aligned BAM files were downloaded from GDC portal (https://portal.gdc.cancer.gov/) for matched paratumor controls (15 additional tissues). The Gene level counts and splice junction counts were derived from AltAnalyze version 2.4.1.2 where junction counts were primarily used for all downstream analyses (gene and splicing). To reduce the impact of sequencing depth differences, we normalized the gene leveland splice junction counts by Count Per Million (CPM). The normalized counts serve as the input matrix **Y**.

We further count the number of samples in each tissue type in which acertain target is detected as expressed (see below). Although intuitively a count of zero should be considered as the cutoff for determining whether a target is detected or not, we observe diverse scenarios where very low normalized counts (less than 1) do not correspond well with the external resources for well-known tissue-specific genes (i.e. NY-ESO-1). We hypothesize choosing a cutoff to filter out the background noise would be beneficial for the purpose of better inference. We determine the cutoff by measuring the concordance between our tissue distribution against the external reference provided by Human Protein Atlas (HPA)^24^. The RNA consensus tissue distribution (RNA consensus tissue gene data) was downloaded from the HPA portal (https://www.proteinatlas.org/about/download). The optimal cutoff can maximize the correspondence (Spearman r correlation) between these two resources (**Supplementary Figure 3**). The thresholded tissue distribution profiles serve as the input matrix **X**.

TCGA Skin Cutaneous Melanoma (SKCM) bulk RNA-Seq BAM files were downloaded from NCI Genomics Data Commons and SRA, following dbGAP authorization.The resultant BAM files were processed using AltAnalyze 2.1.4.2^27^ to obtain the gene count matrix and splice junction matrices. Sashimi plot visualization was generated using ggsashimi package^28^.

### Immunohistochemistry preprocessing

We download non-diseased tissue protein immunohistochemistry annotations (Normal tissue data) from HPA portal (https://www.proteinatlas.org/about/download). The HPA records denote whether a protein product is present or not in a certain tissue type. The annotation contains four levels, namely high, medium, low and not detected. We used a customized script (available at https://github.com/frankligy/BayesTS) to reshape these values into a matrix of the shape (n_protein, 4) where each column corresponds to the number of each annotation level. Since each protein (gene) has a variable number of total slides annotated, we normalized each gene to the average number of slides (P=89 in our data) across all available genes. Finally, we expand the number of labels into categorical values of high, medium, low and not detected are encoded as numerical values 0, 1, 2 and 3, respectively and the total columns amount to P=89. Here the weight vector *w*_*t*_ acts by switching the annotation labels defined by the weight probability (See BayesTS model). The resultant categorical matrix serves as the input matrix **Z**.

### Prior and posterior check

To assure the Bayesian model learns from the observed data in order to adjust for the prior information, we performed prior and posterior checks by simulating samples from both prior and posterior distribution and comparing these against the actual observations. The prior of tumor specificity σ is calculated as the average of 1,000 bootstraps from the randomly initiated Beta(2,2) distribution. The posterior of tumor specificity is calculated as the average of 1,000 bootstraps from the learned Beta distribution parameterized by its alpha and beta parameters in the variational distribution (guide function). For the tissue distribution, we used Poisson distribution with fixed coefficient 25 to simulate a vector of the same length for available tissue (T). For the normalized count, a fixed coefficient 10 was used, as defined by BayesTS model, to simulate out a vector of the same length of all available samples (S). For the protein annotations, we sampled P labels from the underlying categorical distribution with the probabilities determined by the prior or posterior tumor specificities same as the model (See BayesTS model).

### CAR-T therapy targets evaluation

We obtained a list of 71 CAR-T targets that are currently under at least one NCI-registered clinical trials from the CARTSC database^17^ (https://hanlab.tamhsc.edu/CARTSC/#!/). We intersect the 71 targets with the genes that have protein level information available to obtain 55 CAR-T targets for model evaluation.

## RESULTS

### Hierarchical Bayesian modeling to assess tumor specificity

To infer tumor specificity from distinct molecular and distribution-association variables, we developed a hierarchical Bayesian model (BayesTS), without hard-coded thresholds. This model is applied to only non-diseased samples to estimate the probability of a gene/protein being expressed in at least one tissue type. Molecular evidence includes RNA genomic quantification (RNA-Seq) and protein abundance (immunohistochemistry) (**Methods**). The distribution of samples in the reference dataset of normal controls is assessed at a population and tissue level, accounting for varying numbers of replicate controls. The model was developed to allow for flexible weighting of samples associated with distinct tissues in the model, which should be considered as less or not important from the perspective of therapeutic targeting (e.g. testis specific expression in female coded samples).

### BayesTS accurately estimates tumor specificity from multidomain evidence

We trained the BayesTS model using three observations: 1) a normalized RNA count matrix, 2) tissue distribution profiles and 3) protein expression labels (**Methods**). We assure each independent piece of evidence is considered equally in the inference process by scaling their log-probability based on the maximum loss when training them alone (**Methods**). The BayesTS model converges well after around 2000 steps with gradually decreasing ELBO loss, indicating the model successfully updated the weights toward the observed data (**Figure 1B**). To assess whether it can leverage all three pieces of evidence, we project the inferred posterior tumor specificity onto a three-dimensional space where each evidence represents a separate axis. BayesTS is capable of capturing evidence gradients along all three axes to nominate potential targets (**Figure 1C**). To demonstrate the importance of considering multiple types of tumor specificity representations, we performed a sensitivity analysis by removing each modality, one at a time, to evaluate the impact on the final inferred scores. As expected, when protein information was left out of the model, no gradient can be seen along the Z axis, with similar observations along other axes when normalized counts or tissue distributions are not provided to the BayesTS model (**Supplementary Figure 2B, Supplementary Table 1**).

To examine the extent to which proposed BayesTS accurately reflects the tumor specificity of known drug targets we curated 55 CAR-T therapy drug targets that are currently under at least one NCI registered clinical trial (**Methods**). In this set, we find that BayesTS ranked the well-established Cancer Testis Antigens (CTA) MAGEA1 and MAGEA4 as the most tumor-specific targets, due to their restricted expression in testis, which has been used to treat multiple cancers ^29,30^, followed by the highly tumor-specific antigens ALPP ^31^ and CLDN18 (Claudian18.2) ^32^ (**Figure 1D**). We note that not all CAR-T targets have a highly restricted tumor expression profile. Instead, complex optimization steps, such as affinity optimization and drug influx and efflux considerations, are typically used to ensure that the developed therapy only targets tumor cells ^33,34^. Nevertheless, drugs targeting proteins with widespread expression in normal tissues have been reported to cause severe on-target off-tumor toxicity ^35^. Consistent with our sensitivity analysis, when trained using incomplete evidence, the modality that is excluded became less essential for inferred tumor specificity (**Supplementary Figure 2C, Supplementary Table 2**). Importantly, we observe partial correlations between tissue distribution and normalized RNA counts such that even if one piece of evidence is excluded, the overall pattern remains. However, the protein level information is more independent and exhibits a nearly random distribution when it is not included in the BayesTS model (**Supplementary Figure 2C**). This again highlights the need to incorporate multiple pieces of evidence rather than relying on one.

An important validation for any Bayesian model is to inspect whether the model learns from the data and updates the prior belief to better reflect the properties of the observations ^36^. To confirm that the model learns properly, we conducted a prior and posterior check by first generating samples from the underlying prior and posterior distribution of the tumor specificity, and compared these against the actual data distributions. As a lowly expressed target and well-known cancer testis antigens in solid tumors^37^, the tumor specificity of MAGE1A shifts from 0.49 to 0.02 and reflects a clear left-skewed distribution for both the tissue distributions and normalized RNA counts (**Supplementary Figure 4A**). When evaluated on the protein level data, the sampled labels from the posterior have a higher proportion on the “low” label compared to the prior distribution, demonstrating that the model appropriately incorporates information from the data. (**Supplementary Figure 4A**). In contrast, a highly expressed target, UBC, shifts its tumor specificity from prior 0.50 to 0.86 due to the high counts for both its RNA and protein (**Supplementary Figure 4B**). Taken together, we show that BayesTS successfully learns from multiple pieces of evidence and reflects the true distributions for established drug targets.

**Supplementary Figure 2.**
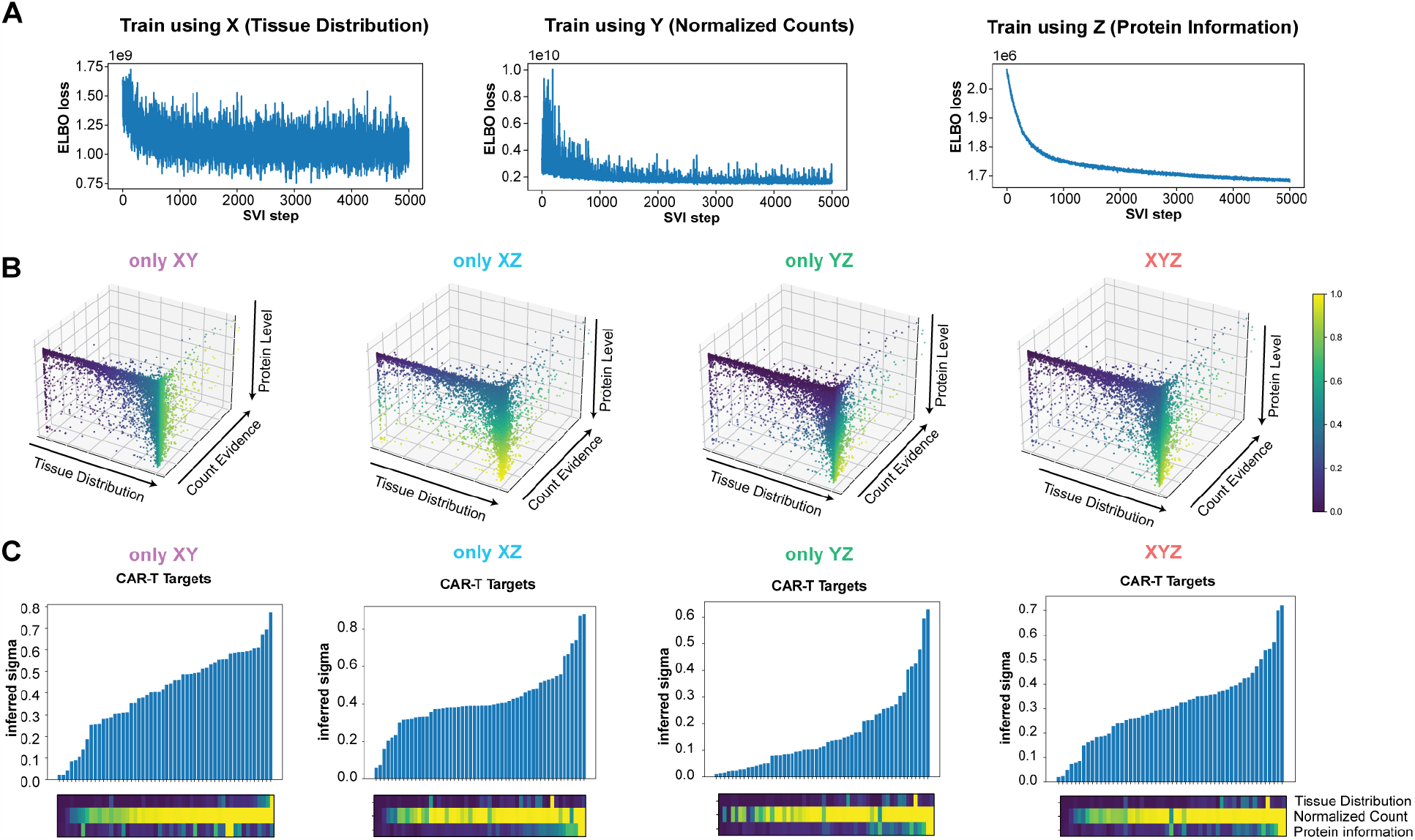
BayesTS incorporates multiple data dimensions for inference. A) The Evidence Lower Bound (ELBO) loss across the training process when only one evidence is provided. B,C) Ablation test on each evidence and the exhaustive combinations of all forms of evidence for all 13,306 protein coding genes (B) and 55 CAR-T targets in clinical trials. C) From left to right are X and Y, X and Z, Y and Z, X, Y and Z.

**Supplementary Figure 3.**
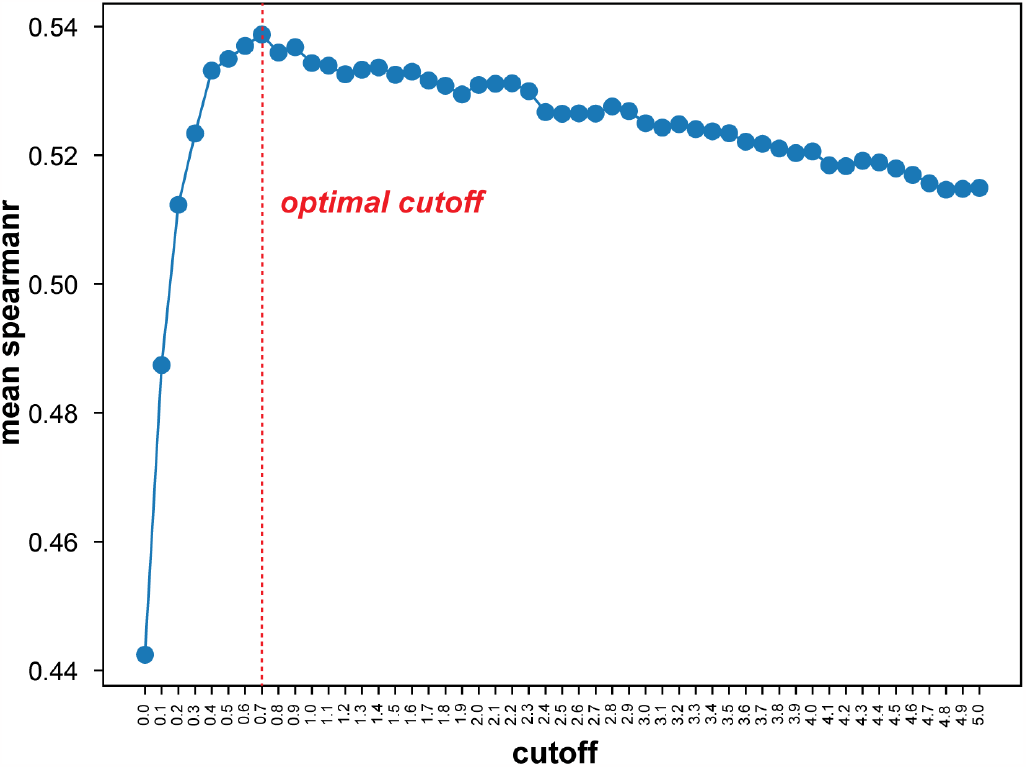
Choosing an optimal cutoff for removing background noise. Spearman correlation was computed between our tissue distribution profiles and external resources. And the optimal threshold is determined by the maximum concordance.

**Supplementary Figure 4.**
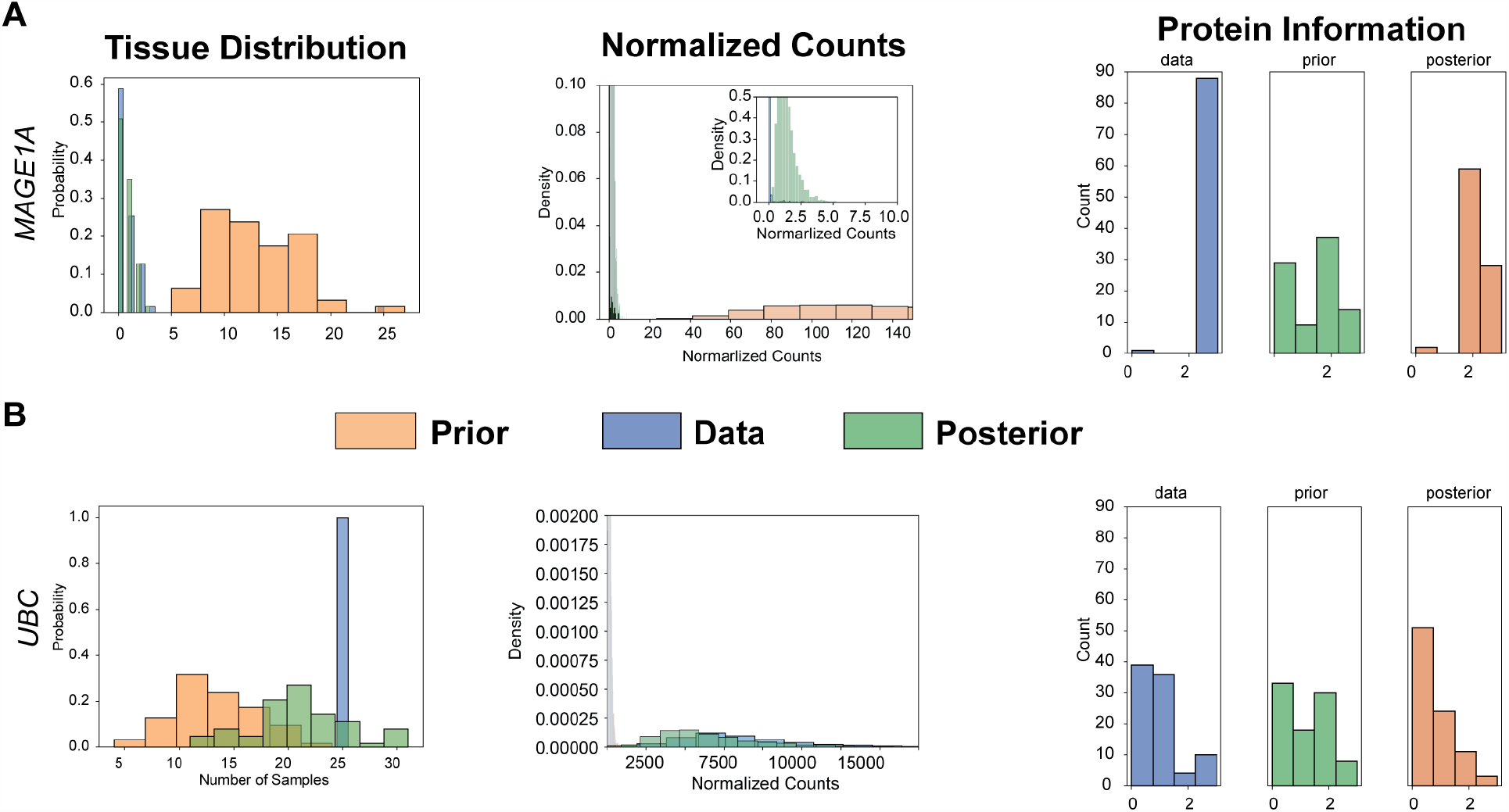
Prior and Posterior check for the BayesTS inference process for two representative targets. A) A lowly expressed target, MAGE1A. B) A highly expressed target, UBC. From left to right, each panel represents one form of evidence, namely, tissue distribution X, normalized RNA count Y and protein-level information Z. Orange represents the prior distribution, blue the observed actual information and green the posterior distribution after the Bayesian variational inference.

### Application of BayesTS to adjust tissue importance and discover novel drug targets

As our first application, we aimed to evaluate the ability of BayesTS to integrat tissue importance information. This feature is particularly valuable as it allows for adaptable adjustment to population-specific cohorts. For example, sex-specific tissues can be selectively down- or up-weighted to accurately reflect the enrolled patient’s tissue type (e.g., skin for melanoma). Additionally, physiologically expendable tissues such as tonsils or appendices, while still significant, can also be preferentially downweighted in defined scenarios. BayesTS empowers users to incorporate prior knowledge of tissue importance to address real-world use cases effectively. We first show the inferred tumor specificity of the Cancer Testis Antigen (CTA) CTAG1B (NY-ESO-1), which decreases from 0.0219 to 0.0114 (Mann Whitney Test p=9.0e-37) when we gradually reduce the the testis importance from 0.5 (default) to 0.2, without violating the overall conclusion that it is still a valid drug target (**Figure 2A**). Next, the tumor specificity for the widely recognized hematological malignancy CAR-T target^38,39^, MS4A1 (CD20), decreases from 0.0814 to 0.0627 (Mann Whitney Test p=2.3e-14) when selectively down-weighting the immune cell producing blood cells and spleen. This approach is particularly relevant because it has been demonstrated to generate a relatively safe clinical response regardless of the presence of remaining immune populations^13^. Furthermore, we additionally down-weighted the tissue importance of appendix and tonsil to 0.1. As expected the tumor specificity continues to decrease for CD20 from 0.0627 to 0.0604 (Mann Whitney Test p=0.07). While the score decrease is relatively minor, this demonstrates the utility of this approach. (**Figure 2A**). The flexibility of BayesTS makes it easy to to be extended in complex clinical trial designs and drug prioritizations.

**Figure 2.**
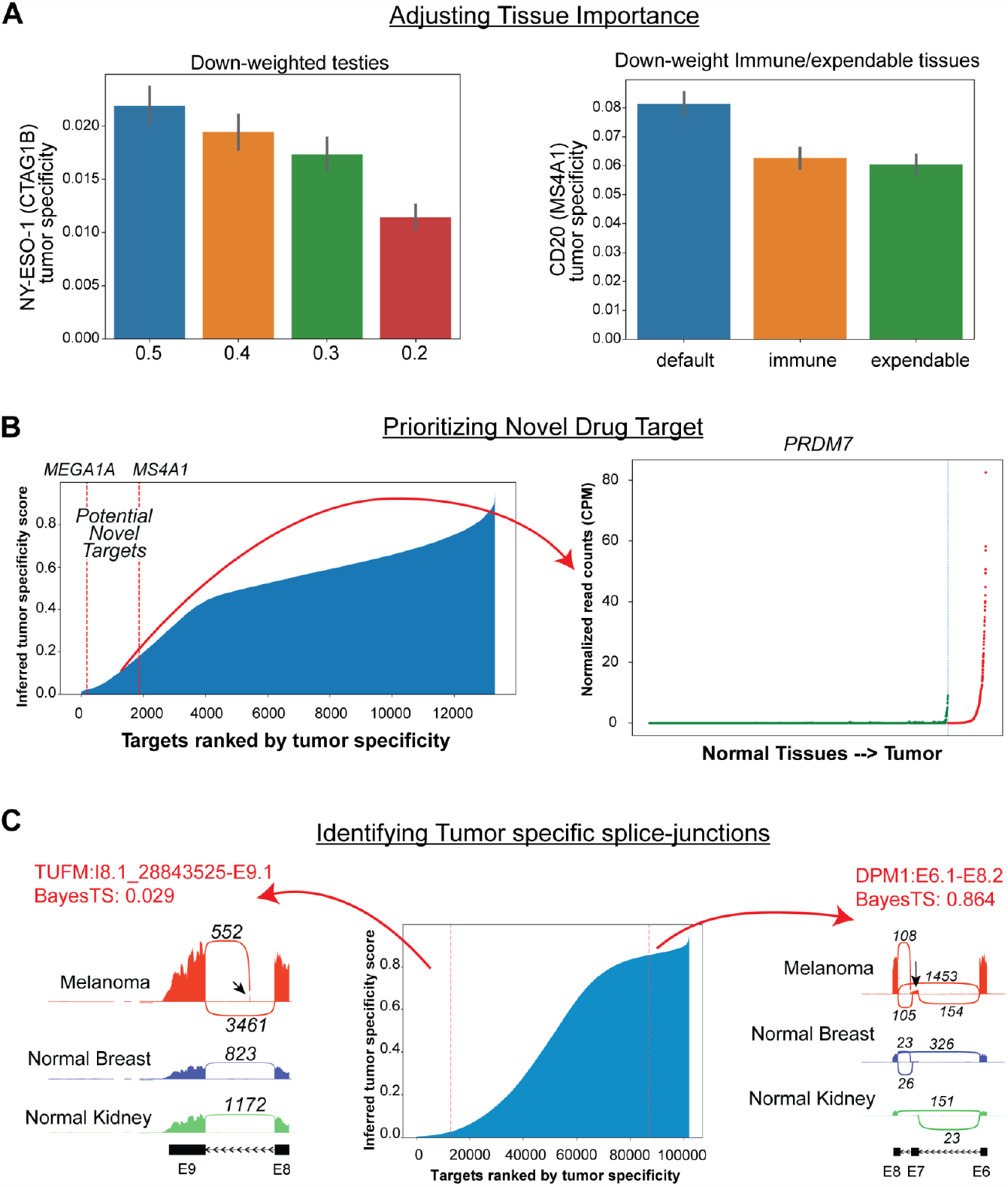
Adjusting tissue importance in the model to identify novel gene targets and unconventional drug targets. A) Down-weighting tissue importance for physiologically irrelevant tissues in the model(left). Here display the inferred changes in sigma (y-axis), after adjusting the tissue importance of testis (left). This is repeated for other select tissues/blood (right), specifically hematological (spleen, blood) to 0.1, and down-weighting of physiologically dispensable tissues (appendix, tonsil) to 0.1. B) Identification of novel drug targets in melanoma from the inferred tumor specificity scores. C) Identification and comparisons of unconventional splice junctions. A tumor-specific novel exon event (left) and an exon-skipping event that are present in normal tissues (right).

As the primary application for a multi-faceted tumor specificity score, BayesTS enables rapid discovery and prioritization of known and novel drug targets. When ranking the tumor specificity of 13,306 genes with protein information in ascending order, targets falling within similar tumor specificity (BayesTS=0.02) as the well-known targets (e.g. MAGE1A, MS4A1) can be selected for further evaluation **(Figure 2B)**. To identify novel targets, we analyzed TCGA Skin Cutaneous Melanoma (SKCM) with 472 patients in total and compared the gene expression differences of nominated drug targets based on inferred tumor specificity score. First, we found multipleMAGE family proteins (e.g. MAGEC2, MAGEB2, MAGEC1, MAGEB6) with a similar gene expression profile to those inclinical trials (MAGEA1, MAGEA4), which have been reported to be tumor specific ^40–42^. Second, we find numerous Pregnancy-specific glycoprotein (PSG) genes which fall into the broader category of carcinoembryonic antigens (**Supplementary Figure 6A**). While the association of placenta trophoblasts and tumor have been previously suggested ^43^, these targets have not been included in active CAR-T clinical trials, hence, these may represent novel unexploited targets for melanoma treatment. Asides from these previously reported potential antigens, we nominated a histone methyltransferases gene (PRDM7) that is restricted in its expression to testis, but that is otherwise uniquely expressed in melanoma patients. PRDM7 is responsible for regulating DNA methylation and accessibility (H3K4me2 to H3K4me3), and hence could potentially play a role in tumor homeostasis (**Figure 2B**). In addition, these analyses highlight a number of novel targets that are expressed in melanoma (RNA-Seq) but have evidence of a lack of expression in healthy tissues from GTEx, TCGA controls and HPA. These include the N-terminal acetyltransferases (NAA11), a suppressor of circadian rhythm (PASD1), the suppressor of microRNA biogenesis (LIN28) and the transcriptional repressor (SSX4) (**Supplementary Figure 6**). Hence, such proteins represent intriguing high-value targets for validation in independent control tissue datasets.

**Supplementary Figure 5.**
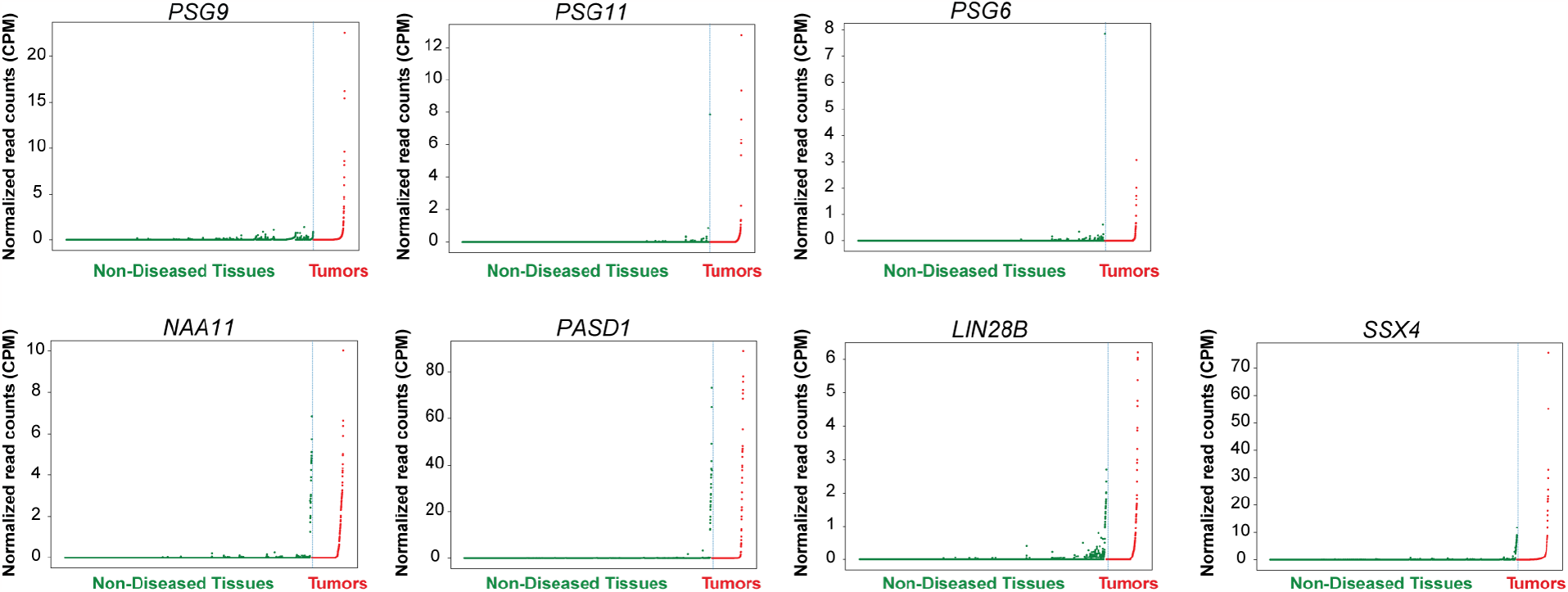
Identified novel drug targets using BayesTS. Tissue-specificity RNA expression plots for control tissues (green) and melanoma patient samples (red), where each point is an independent sample (blue lines separating normal and tumor tissue samples, organized by tissue type). Examples are shown for Pregnancy-specific glycoproteins (PSG), namely PSG6, PSG9 and PSG11 and other potentially novel targets for therapy.

**Supplementary Figure 6.**
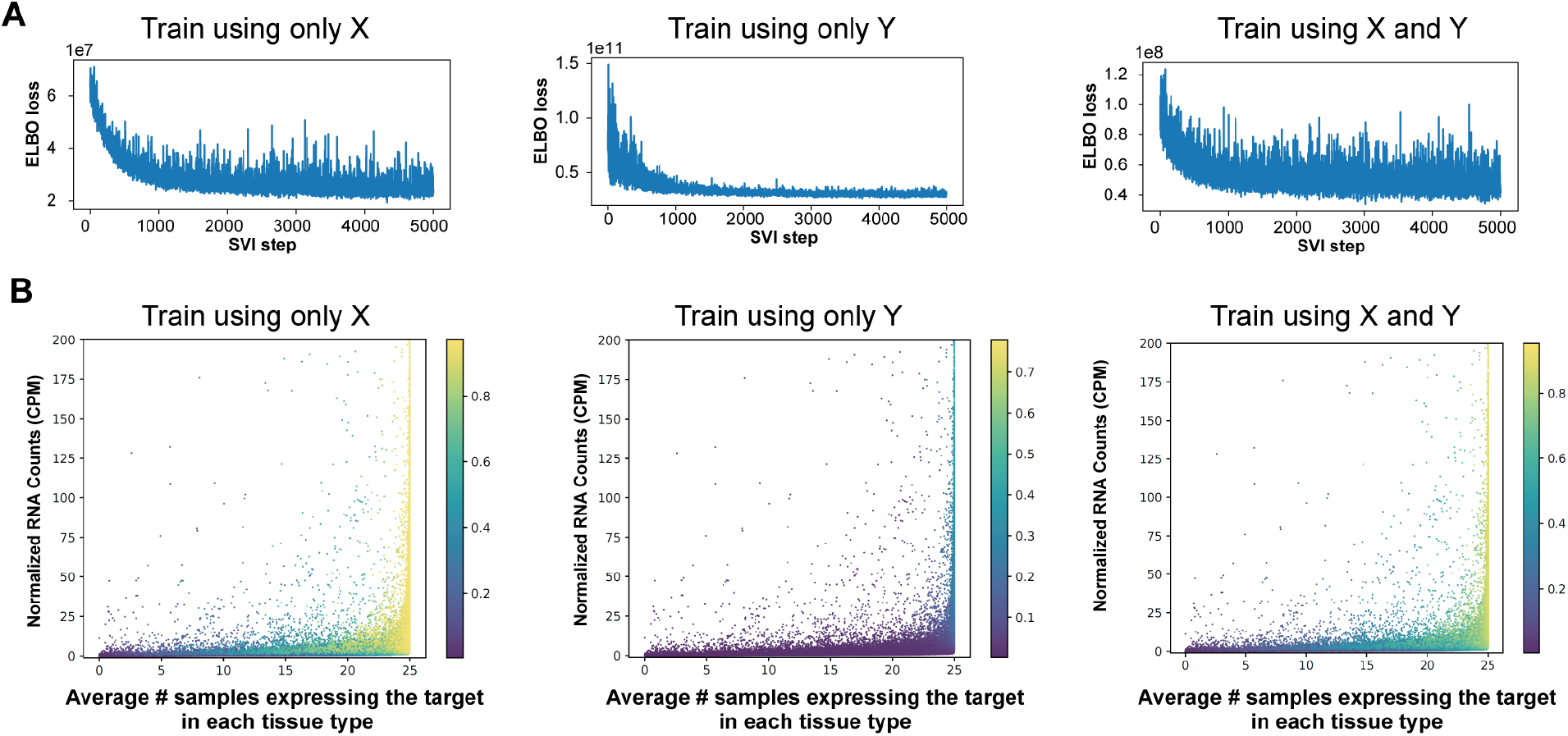
Training loss and validation of BayesTS on splice junction targets. A) The ELBO loss of training tissue distribution (X) or normalized RNA count (Y) separately and together (right). B) The inferred tumor specificity scores projected on both X and Y axis in three different scenarios.

Finally, to demonstrate thatBayesTS can be readily applied to mechanistically distinct molecular targets We applied this model to a large datasets of tumor-detected alternative splicing events (melanoma). Tumor-specific splicing events have been increasingly recognized as ubiquitous hallmarks of diverse malignancies ^44^, that are often associated with clinical outcome^45^. Such targets include splicing neoantigens, which are highly specific exon-exon or exon-intron junctions that encode for immunogenic MHC presented peptides, representing new targets for cancer vaccine development ^12,46^. We analyzed 472 TCGA Skin Cutaneous Melanoma (SKCM) patients and quantified individual sample splice junction expression. Similar to our gene-focused pipeline, we trained the BayesTS model using only tissue distributions and normalized RNA counts for GTEx and TCGA control splice-junctions. Here, we were not able to supply associated protein quantifications for these junctions, which require highly sensitive and specific junction-level mass spectrometry profiles. First, we re-derived the scaling factors by training each evidence (**Supplementary Figure 6A**). The scaling factors ensure each evidence contributes equally to the inference. When combined, BayesTS learns toward the observed data, while leaving one modality out fell short of capturing the corresponding information (**Supplementary Figure 6A,B**). We highlight two splice junctions with opposite tumor specificity verified by SashimiPlot visualization. These include, observe a novel exon of gene TUFM (I8.1_28843525-E9.1) that is highly specific to melanoma patients (BayesTS score = 0.029), whereas a exon skipping event DPM1 (E6.1-E8.2) are broadly present in normal tissues (BayesTS score = 0.864) (**Figure 2C, Supplementary Table 3**). Hence, BayesTS is broadly applicable to different molecular modalities and molecular specificity contexts.

## DISCUSSION

Accurately predicting tumor specificity is a crucial step in the design of new therapeutic strategies for diseases such as cancer to improve target prioritization and preclinical evaluation. We proposed a new index, BayesTS, to simultaneously connect multiple representations of tumor specificity including both RNA and protein evidence. Using a large dataset of normal tissue samples across over 3,500 samples spanning 63 tissues, we showed that BayesTS can prioritize well-established drug targets while deemphasizing those that induce toxicity. Our approach has several advantages over existing approaches, which rely on ad-hoc thresholds and manual inspection. Further, BayesTS can be easily extended to other modality such as epigenetic^47^ and Post-Translational Modification (PTM) profiles^48^.

We note that there are potential limitations of our approach, both in terms of our overall assumptions and potential Type 2 errors. First, our approach is restricted to the analysis of normal control tissues in which BayesTS assesses the probability of absence of a signal. While tumor data is needed to identify highly expressed tumor antigens, BayesTS represents a critical filter for such analyses, which we find reduces the search space from thousands of targets to dozens or hundreds. A second caveat is that this approach assumes the same detection biases exist in the control and disease tissues evaluated. If significant differences in the RNA or protein quantification exist between the control and disease datasets (e.g., batch effects, library generation, sequencing depth), this could result in both false positive and false negative predictions. Finally, as our approach has been applied to bulk RNA-Seq and protein quantification data, it is likely to blind to extremely rare biologically crucial cell-type gene expression programs in healthy cells, which express otherwise predicted tumor specific molecules. This challenge can be overcome through extension of this approach to large single-cell and disease cohorts with different sample and cell-population biases, that can be accounted for automatically in the underlying model. Furthermore, Multi-view deep neural network based representation learning may be capable of capturing complex interactions amongst different forms of evidence better than explicit Bayesian modeling ^49^. However, we expect our Bayesian strategy to be applicable to diverse disease specificity challenges, exploiting both tissue and cell-level candidate target abundance. To facilitate its immediate use for diverse clinical applications, we provide BayesTS as a queryable database of tumor specificity scores.

## ACKNOWLEDGMENTS

This work was supported by generous funding from Cincinnati Children’s Hospital Research Foundation and the National Institutes of Health (R01CA226802 to N.S.).

## CONTRIBUTIONS

G.L. conceived the method, wrote the software and performed all bioinformatics analyses. A.B. assisted with data acquisition, processing and quality control analyses. G.L and N.S. wrote the manuscript.

## Notes

https://github.com/frankligy/BayesTS

https://www.synapse.org/#!Synapse:syn51170082/files/

### Competing Interest Statement

The authors have declared no competing interest.

